# Conscientiousness associated with efficiency of the salience/ventral attention network: Replication in three samples using individualized parcellation

**DOI:** 10.1101/2022.06.07.495168

**Authors:** Tyler A. Sassenberg, Philip C. Burton, Laetitia Mwilambwe-Tshilobo, Rex E. Jung, Aldo Rustichini, R. Nathan Spreng, Colin G. DeYoung

## Abstract

Previous research in the field of personality neuroscience has identified associations of conscientiousness and related constructs like impulsivity and self-control with structural and functional properties of particular regions in the prefrontal cortex (PFC) and insula. Network-based conceptions of brain function suggest that these regions probably belong to a single large-scale network, labeled the salience/ventral attention network (SVAN). The current study tested associations between conscientiousness and resting-state functional connectivity in this network using two community samples (*N =* 244 and 239) and data from the Human Connectome Project (*N* = 1000). Individualized parcellation was used to improve the accuracy of functional localization and to facilitate replication. Functional connectivity was measured using an index of network efficiency, a graph theoretical measure quantifying the capacity for parallel information transfer within a network. Efficiency of a set of parcels in the SVAN was significantly associated with conscientiousness in all samples. Findings are consistent with a theory of conscientiousness as a function of variation in neural networks underlying effective prioritization of goals.

---

Within the Five Factor Model of personality, conscientiousness describes the shared variance among traits reflecting tendencies to follow rules and prioritize non-immediate goals (DeYoung, 2015). Individuals scoring high in conscientiousness are inclined toward fastidiousness, hard work, future planning, and are skilled at self-regulating and avoiding impulsivity. Conscientiousness predicts a wide variety of influential life outcomes, including health-promoting behaviors, longevity, quality of familial and intergenerational relationships, academic success, workplace performance, and career success (Ozer & Benet-Martinez, 2006; Roberts et al., 2014). In predicting career and academic outcomes, it is second only to intelligence (Wilmot & Ones, 2019; Higgins et al., 2007). Further, conscientiousness represents the opposite pole of the dimension of psychopathology known as disinhibition, which is associated with externalizing disorders such as attention deficit/hyperactivity disorder (ADHD) and substance use disorders (Widiger et al., 2019). Despite the clear importance of conscientiousness for human life, relatively little is established regarding its underlying neural substrate (DeYoung et al., 2021).

Based on the existing research on neural correlates of conscientiousness and impulsivity, Allen and DeYoung (2017) theorized that conscientiousness is in part a function of a particular broad neural network that has been identified in large studies of functional connectivity (Schaefer et al., 2018; Uddin et al., 2019; Yeo et al., 2011). This network includes core neuroanatomical regions, the anterior insula and dorsal anterior cingulate cortex (dACC), and has been studied under the labels of “salience,” “ventral attention,” and “cingulo-opercular” networks, (Dosenbach et al., 2007; Fox et al., 2006; Seeley et al., 2007; Uddin et al., 2019). Here we refer to it as the salience/ventral attention network (SVAN). Considering the known functional properties of these networks, Rueter et al. (2018) suggested that, as a whole, the SVAN might be considered a goal priority network, responsible for prioritizing goals effectively given situational affordances and directing attention away from distractions and toward goal-relevant stimuli. In addition to the insula and dACC, the SVAN includes nodes in lateral PFC, including dorsolateral PFC, inferior parietal cortex (operculum) and temporoparietal junction (TPJ) (Yeo et al., 2011; Uddin et al, 2019).

Rueter et al. (2018) attempted to perform the first direct test of the hypothesis that functional connectivity within the SVAN is positively associated with conscientiousness. However, limitations in their methods rendered it unclear to what extent their findings would generalize to other samples and how accurately they captured variance associated with the SVAN specifically. They used networks derived from independent components analysis (ICA) that did not align perfectly with the borders of the SVAN found in standard atlases. Additionally, ICA solutions are unique to the sample in which the ICA was conducted, which impedes replication. In the present research, we rely on the atlases of Yeo et al. (2011) and Schaefer et al. (2018) to define the SVAN. This allowed us to specifically test the SVAN hypothesis and to reproducibly assess the same functional neuroanatomy across multiple samples.

A major challenge when testing hypotheses regarding individual differences in networks from standard atlases is that these networks are not in the same spatial locations for everyone, relative to anatomical landmarks (Chong et al., 2017; Gordon et al., 2017; Kong et al., 2018). Thus, if one simply overlays a standard network atlas on each individual’s structurally aligned data, the boundaries will be poorly estimated for everyone. In order to better estimate regional boundaries, those boundaries must be adjusted for each subject. To accomplish that adjustment, we employed a relatively new technique to accomplish the individualization of network locations, based on an iterative Bayesian process, known as group prior individualized parcellation (GPIP; Chong et al., 2017). In GPIP (and similar procedures, e.g. Kong et al., 2021), one begins with the boundaries of a standard atlas, either of distributed networks or of smaller contiguous parcels. Then patterns of covariance in fMRI data for each subject are used to adjust the boundaries of the atlas to optimize them to that subject’s unique pattern of functional connectivity. Thus, all networks or parcels are present in each subject and retain their identity, but their locations are individualized.

Individualized parcellation inherently produces greater within-parcel covariance, and this seems to be a good marker of true functional coherence. Compared to non-individualized atlases, individualized parcellations correspond better to regions of task-related activation, which conform more closely to functional organization than to anatomical location (Chong et al., 2017). It also improves prediction of a wide range of individual differences (Anderson et al., 2021; Kong et al., 2021; Setton et al., 2022). Coupled with the fact that individual parcels from standard atlases can easily be compared across subjects and samples, these findings suggest that individualized parcellation should become standard practice in neuroimaging research on individual differences (DeYoung et al., in press).

We tested the theorized association of conscientiousness with connectivity in the SVAN, using this approach in the original sample studied by Rueter et al. (2018) as well as two additional independent samples, including data from the Human Connectome Project (HCP). We used an atlas by Schaefer et al. (2018) with 400 parcels that align very closely with the canonical networks identified by Yeo et al. (2011). Our hypothesis was that the functional connectivity among parcels in the SVAN would be positively associated with conscientiousness. We quantified functional connectivity using a measure of network efficiency that reflects the capacity for parallel information transfer within a network and that has been identified as a reliable metric for characterizing functional network integration (Bullmore & Sporns, 2012; Deuker et al., 2009; Jiang et al., in review). To avoid assuming that every parcel in the network must be equally important to conscientiousness, we also examined subsets of parcels within the SVAN and its subnetworks.

## Methods

### Sample 1

#### Participants

A total of 306 right-handed participants completed a single resting-state fMRI session as part of a larger study. Participants were recruited through online advertisements and fliers posted in public areas in the metro region around Minneapolis and St. Paul, Minnesota. Exclusion criteria include fMRI contraindications, a diagnosis of neurological or severe psychiatric conditions, or substantial behavioral dysfunction attributable to drug or alcohol use. Following recruitment, additional participants were excluded due to incomplete or poor quality fMRI data, incomplete behavioral data, poor FreeSurfer surface alignment, and excessive head movement during the scan, as identified through multiple data preprocessing pipelines applied in subsequent analyses. A total of 244 subjects were retained (121 females) ranging from 20 to 40 years old (*M* = 25.9, *SD* = 4.7). Protocols used in this study were approved by the University of Minnesota Twin Cities institutional review board, and all participants provided written informed consent.

#### Personality Measures

Participants in the present study completed two personality questionnaires, the Big Five Aspect Scales (BFAS; DeYoung et al., 2007), and the Big Five Inventory (BFI; John et al., 2008). The BFAS comprises 100 items on a 5-point Likert scale ranging from 1 (*strongly disagree*) to 5 (*strongly agree*). The questionnaire measures two lower-order aspects for each of the Big Five, with 10 items per aspect. These aspect scores can be averaged to create 20-item domain level Big Five scores. The BFI consists of a total of 44 items measuring domain level Big Five factors on a 5-point Likert scale ranging from 1 (*strongly disagree*) to 5 (*strongly agree*). In addition to self-report measures, peer-report measures were obtained by providing participants with 3 packets with instructions to have both questionnaires completed by individuals who knew the participant well. At least one peer report was available for 182 participants, and multiple peer-reports for a given subject were averaged to create a single peer-report score, when applicable. Scale scores for Conscientiousness across the BFAS and BFI were averaged to create a composite variable, and self and peer-report measures were averaged when applicable.

#### Intelligence

All participants completed a subset of the Wechsler Adult Intelligence Scale – Fourth Edition (WAIS-IV; Wechsler, 2008), the Block Design, Matrix Reasoning, Vocabulary, and Similarities tests. Although the present study selected these tests from the full WAIS-IV, these four tests are identical to those used in the shorter alternative, the Wechsler Abbreviated Scale of Intelligence (WASI), and provide reliable estimates of full-scale IQ (Wechsler, 2011).

#### fMRI Data Acquisition and Preprocessing

fMRI data were acquired using a 3T Siemens Trio Scanner at the Center for Magnetic Resonance Research at the University of Minnesota Twin Cities. High-resolution T1-weighted MPRAGE images with the following parameters were acquired for anatomical surface registration for each participant: voxel dimensions = 1 x 1 x 1 mm^3^; repetition time (TR) = 1.9 s; echo time (TE) = .29 ms; flip angle = 9°. Functional echo-planar images were acquired with the following parameters: 35 coronal slices; TR = 2 s; TE = 28 ms; flip angle = 80°; voxel dimensions = 3.5 x 3.5 x 3.5 mm.

Results included in this manuscript come from preprocessing performed using *fMRIPrep* version 20.2.1 (Esteban et al., 2018; RRID:SCR_016216), a *Nipype* (Gorgolewski et al., 2011; RRID:SCR_002502) based tool. Each T1w (T1-weighted) volume was corrected for INU (intensity non-uniformity) using N4BiasFieldCorrection v2.1.0 (Tustison et al., 2010) and skull-stripped using antsBrainExtraction.sh v2.1.0 (using the OASIS template). Spatial normalization to the ICBM 152 Nonlinear Asymmetrical template version 2009c (Fonov et al., 2009; RRID:SCR_008796) was performed through nonlinear registration with the antsRegistration tool of ANTs v2.1.0 0 (Avants et al., 2008; RRID:SCR_004757), using brain-extracted versions of both T1w volume and template. Brain tissue segmentation of cerebrospinal fluid (CSF), white-matter (WM) and gray-matter (GM) was performed on the brain-extracted T1w using fast (FSL v5.0.9; Zhang et al., 2001; RRID:SCR_002823).

Functional data was slice time corrected using 3dTshift from AFNI v16.2.07 (Cox, 1996; RRID:SCR_005927) and motion corrected using mcflirt (FSL v5.0.9; Jenkinson et al., 2002). This was followed by co-registration to the corresponding T1w using boundary-based registration (Greve & Fischl, 2009) with nine degrees of freedom, using flirt (FSL). Motion correcting transformations, BOLD-to-T1w transformation and T1w-to-template (MNI) warp were concatenated and applied in a single step using antsApplyTransforms (ANTs v2.1.0) using Lanczos interpolation.

Physiological noise regressors were extracted applying CompCor (Behzadi et al., 2007). Principal components were estimated for the two CompCor variants: temporal (tCompCor) and anatomical (aCompCor). A mask to exclude signal with cortical origin was obtained by eroding the brain mask, ensuring it only contained subcortical structures. Six tCompCor components were then calculated including only the top 5% variable voxels within that subcortical mask. For aCompCor, six components were calculated within the intersection of the subcortical mask and the union of CSF and WM masks calculated in T1w space, after their projection to the native space of each functional run. Frame-wise displacement (Power et al., 2014) was calculated for each functional run using the implementation of *Nipype*. ICA-based Automatic Removal Of Motion Artifacts (AROMA) was used to generate aggressive noise regressors as well as to create a variant of data that is non-aggressively denoised (Pruim et al., 2015). Participants exhibiting a relative mean framewise displacement greater than 0.5 mm, a mean standardized derivative of RMS variance over voxels (DVARS) greater than 1.5, or any single occurrence of a coordinate displacement greater than 2.75 mm were excluded from subsequent analyses to avoid biased effects in associations with measures of functional connectivity (Power et al., 2012; Power et al., 2014).

### Sample 2

#### Participants

A total of 260 participants were recruited through postings throughout the University of New Mexico, nearby high schools, and various professional STEM businesses from communities surrounding Albuquerque, New Mexico. Participants were excluded on the basis of neurological and psychological disorders, fMRI contraindications, incomplete behavioral data, incidental fMRI findings, poor FreeSurfer surface registration, and excessive head motion identified through data preprocessing. A total of 239 participants were retained for the present study (117 females) ranging from 16 to 38 years old (*M* = 22, *SD* = 3.9). All participants provided written informed consent, and all procedures in this study were approved by the University of New Mexico institutional review board.

#### Personality Measures

All participants included in the present study completed one of two personality questionnaires: the BFAS, or the NEO Five-Factor Inventory (FFI). The NEO-FFI represents a subset of the full NEO Personality Inventory, Revised (NEO PI-R; Costa & McCrae, 1992), consisting of 12 items per factor. Scale scores were calculated as item averages, using a five-point Likert scale ranging from 0 (*strongly disagree*) to 4 (*strongly agree*). A total of 59 participants completed the NEO-FFI, and 180 participants completed the BFAS. To account for differences in scale means, scores were centered by subtracting the mean of their respective scale. The present study utilized Conscientiousness scores from all participants.

#### Intelligence

All participants were administered the set of four tasks from the WASI (Wechsler, 2011). Full-scale IQ was estimated from performance on the Vocabulary, Similarities, Matrix Reasoning, and Block Design subtests.

#### fMRI Data Acquisition and Preprocessing

Using a 3T Siemens Prisma scanner, resting-state functional echo-planar images were acquired with the following parameters: 32 coronal slices; TR = 275 ms; TE = 30 ms; flip angle = 34°; multiband acceleration factor = 8; voxel dimensions = 3.5 x 3.5 x 3.5 mm, pixel bandwidth = 1736 Hz. High-resolution T1- weighted MPRAGE images were acquired for anatomical surface registration through a 5 echo sequence with the following parameters: TR = 25.3 s; TE = 1.64 ms, 3.5 ms, 5.36 ms, 7.22 ms, 9.08 ms; flip angle = 7°; voxel dimensions = 1 x 1 x 1 mm^3^. Results included in this manuscript come from preprocessing performed using *fMRIPrep* version 20.2.1 using the same specifications described for Sample 1.

## Sample 3

### Participants

A total of 1000 participants (533 females) were selected from the WU-Minn Consortium of the Human Connectome Project (Van Essen et al., 2012). Participants were initially excluded on the basis of a history of severe psychiatric, neurological, or medical disorders. Among the participants of the 1200 young adult sample, additional participants were excluded due to missing personality and intelligence task data, and missing resting-state fMRI scan data. Participants’ ages ranged from 22 to 37 years old (*M* = 28.7, *SD* = 3.7). All participants provided informed consent, and all study protocols were approved by the Institutional Review Board of Washington University in St. Louis. Details of the informed consent procedure are provided by Van Essen et al. (2013).

### Personality Measures

Participants completed the NEO-FFI to assess trait Conscientiousness. Scale scores were assessed using a five-point Likert scale ranging from 0 (*strongly disagree*) to 4 (*strongly agree*).

### Intelligence

Intelligence was assessed as a composite of performance measures from a set of three tests from the NIH Toolbox (Heaton et al., 2014) and Penn Computerized Neurocognitive Battery (Moore et al., 2015). Performance metrics on the Matrix Reasoning, Picture Vocabulary, and List Sorting tasks were averaged to create an estimate of intelligence for analyses.

### fMRI Data Acquisition and Preprocessing

fMRI data were acquired using a customized 3T Siemens Skyra scanner for all participants at Washington University in St. Louis. The present study used a single left-to-right phase encoded resting-state scan acquired using the following parameters: 72 axial slices; TR = 0.4 s; TE = 33 ms; flip angle = 52°; multiband acceleration factor = 8; voxel dimensions = 2 x 2 x 2 mm^3^; pixel bandwidth = 2,290 Hz. Additionally, high-resolution T1-weighted MPRAGE structural images were acquired for anatomical surface registration with the following parameters: TR = 24 s; TE = 2.14 ms; flip angle = 8°; voxel dimensions = 0.7 x 0.7 x 0.7 mm^3^. Resting-state scans were preprocessed using the HCP minimal preprocessing pipeline and motion artifacts were removed using ICA-FIX (Burgess et al., 2016). Relative mean framewise displacement was also computed to be included as a covariate in subsequent analyses. Features of the HCP minimal preprocessing pipeline are described in greater detail in previous literature (Glasser et al., 2013; Ugurbil et al., 2013).

### Group Prior Individualized Parcellation

Functional connectivity networks were identified using an individualized cortical parcellation approach. For participants in all three samples, ICA-denoised resting-state fMRI scans in subject-native space were first resampled to a common cortical surface mesh (Dale et al., 1999), and the BOLD timeseries at each vertex was normalized to zero mean and unit variance. The resulting subject surface data were then initialized using a pre-defined group atlas with 400 functionally distinct regions (Schaefer et al., 2018) mapped to the 17-network atlas defined by Yeo et al. (2011). A Bayesian algorithm was applied to iteratively adjust parcel boundaries according to each participant’s unique patterns of functional connectivity (Chong et al., 2017). In order to produce stable modifications of parcel boundaries, this algorithm utilized 20 iterations to ensure that all participants had no more than one vertex changing its parcel label on the final iteration. Through this process, each participant acquired a unique modification of a standard group-level atlas after the final iteration such that the boundaries of the parcels of the initial atlas optimally reflected each individual’s patterns of resting-state functional connectivity.

For the present analyses, the full SVAN was defined by 51 functional parcels, 34 of which correspond to subnetwork A, and 17 of which correspond to subnetwork B, in the 17- network atlas of Yeo et al. (2011). The full SVAN was included in the present analyses to account for the potential role of connections between parcels of subnetworks A and B, rather than focusing only connections within each subnetwork separately. (The two subnetworks fractionate the larger network labeled “ventral attention network” in Yeo et al.’s (2011) 7- network atlas.) The parcels of the SVAN are illustrated using the group-level Schaefer atlas (Schaefer et al., 2018) in Figure 1.

**Figure 1.**
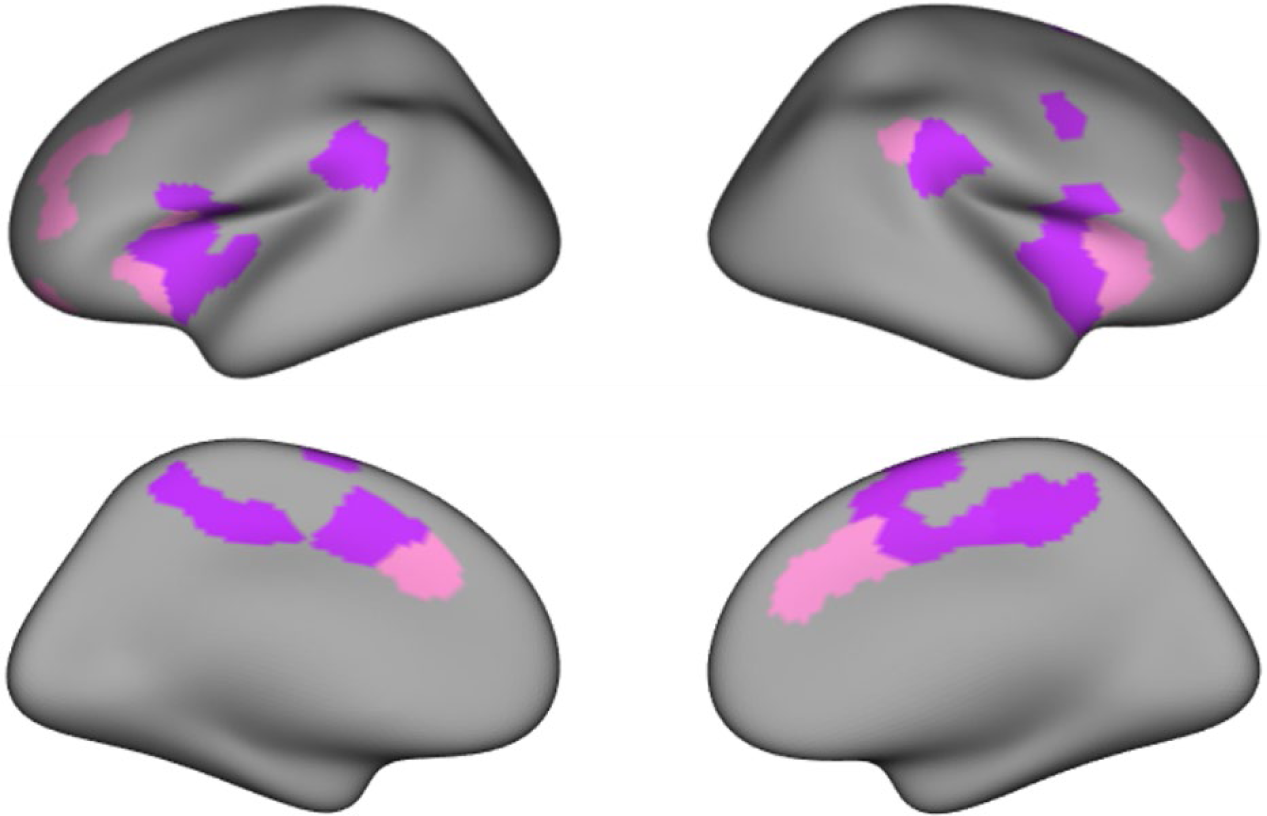
Parcels in the salience/ventral attention network (SVAN). Colors correspond to 2 of the 17 functional networks described by Yeo et al. (2011). Purple = SVAN subnetwork A, Pink = SVAN subnetwork B

### Orthogonal Minimum Spanning Trees and Functional Connectivity

For each sample, individualized parcel average timeseries were correlated and Fisher r- to-z transformed to create a 400 × 400 functional connectivity matrix for each participant. Using these matrices, smaller separate matrices were constructed for correlations among parcels in the SVAN, subnetwork A, and subnetwork B. To quantify the connectivity properties of each matrix, a graph-theory approach was used, in which each parcel is considered a node in the graph and each correlation represents a potential connection, or edge, between nodes.

We applied a data-driven topological threshold to the individual functional connectivity matrices using orthogonal minimum spanning trees (OMSTs; Dimitriadis et al., 2017). The OMST approach was used to screen out many low correlations and avoid modeling noise that may be present in functional connectivity measures, while also avoiding the imposition of arbitrary graph thresholds that incur methodological biases. OMSTs rely on an iterative procedure to filter functional connectivity networks to optimize the trade-off between efficiency within the network and the wiring cost, where wiring cost refers to the ratio of the sum of the weights of the filtered graph to that of the unfiltered graph. In this manner, connections between participants’ OMSTs using an identical set of nodes can vary substantially on account of unique individuals’ patterns of brain organization, while still ensuring a fully connected graph, a property generally exhibited by unfiltered weighted graphs. We emphasize the importance of utilizing a fully connected graph for each network of interest to reflect the notion that these parcels demonstrate functional interactions within canonical networks described by the group-level cortical atlas. Despite the application of a degree of sparsity to the functional connectivity matrices using OMSTs, this thresholding procedure has been demonstrated to produce (1) graphs characterized by similar sensitivity to perturbations in connection strength and density compared to unfiltered graphs (Tewarie et al., 2015), (2) higher performance compared to conventionally filtered graphs in tests of subject-specific graph recognition (Dimitriadis et al., 2017), and (3) greater test-retest reliability across scans compared to graphs with conventional thresholds (Jiang et al., in review).

We computed efficiency of OMST-filtered graphs as an index of network-wide functional connectivity. Efficiency describes the average inverse shortest path length within the network (Rubinov & Sporns, 2010). In the present analyses, efficiency reflects coherence among sets of parcels as a function of correlations between average parcel timeseries. The choice of this particular graph theoretical measure is informed by research demonstrating its reliability as a measure of functional network integration (Bullmore & Sporns, 2012; Deuker et al., 2009; Jiang et al., in review).

### Analysis

We used two main analytic strategies. First, we examined associations between conscientiousness and efficiency among all parcels within each network of interest. Second, to address the potential influence of noise modeled in pairwise functional connectivity metrics and to constrain the search space of influential edges across multiple samples, we applied a permutation-based feature selection approach to identify potentially relevant subsets of parcels within each network. This approach is inspired by the data-driven, machine-learning method called Connectome-Based Predictive Modeling (CPM; Finn et al., 2015). CPM uses cross-validation to identify unique brain-behavior associations using functional connectivity data and has been effective in identifying significant and reliable associations across large samples (Rosenberg et al., 2015; Wang et al., 2021). However, since CPM is primarily a data-driven procedure, using this method across multiple samples introduces the likely possibility of identifying different configurations of edges in each, which is not suitable for the goal of identifying replicable groups of parcels. To take advantage of CPM’s capacity to summarize broad patterns of functional connectivity, we used a variation of the feature selection criteria described in the CPM procedure to identify a consistent set of parcels whose associations with conscientiousness could be tested across multiple samples.

For tests of associations between conscientiousness and subsets the networks of interest (i.e., SVAN and its subnetworks A and B), we randomly divided participants in each sample into deciles. Using nine of the ten deciles, partial correlations were computed between each edge in OMST-filtered functional connectivity matrices and conscientiousness, controlling for the effects of head motion using relative mean framewise displacement. This procedure was repeated ten times, holding out each decile once. In each iteration, a binarized matrix of the graph was created denoting edges positively associated with conscientiousness at *p* < .05 (uncorrected). Across the ten iterations, these matrices were combined to represent the frequency of iterations in which edges demonstrated nominally significant associations with conscientiousness. The maximum value of each row in this matrix was used to signify the robustness to sampling variability of associations between conscientiousness and the connectivity of each parcel. For each sample, parcels were selected for further analyses if they contained an edge significantly positively associated with conscientiousness in at least 8 iterations. The division of each sample into deciles was motivated by the desire to reduce the computational intensity of the procedure while also providing each parcel with an index of its association with conscientiousness in line with conventional thresholds used in CPM (Shen et al., 2017). This workflow is illustrated in Figure 2.

**Figure 2.**
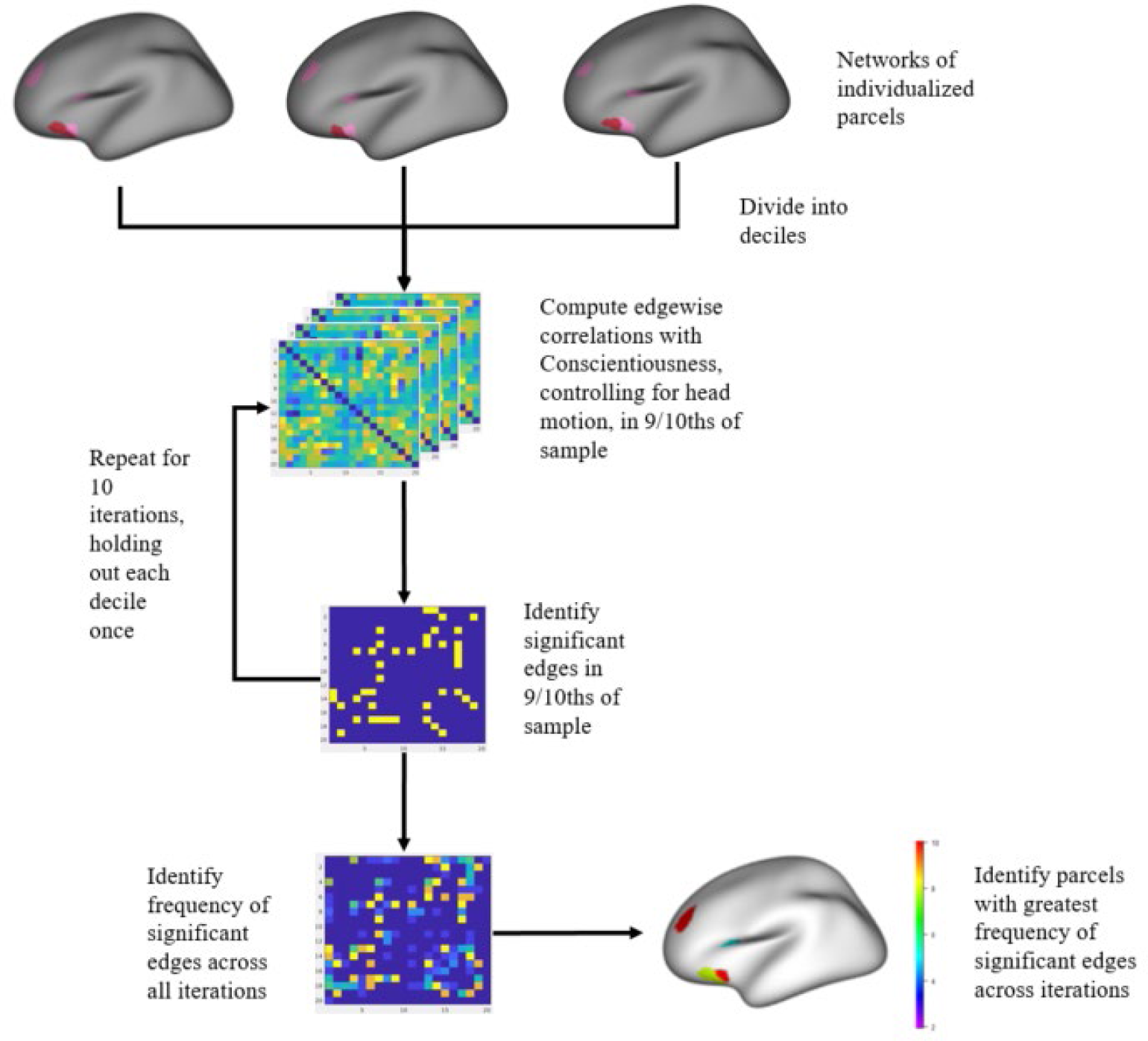
Workflow of method for identifying subnetworks in each sample. This procedure was conducted in all samples, but parcels in Sample 3 did not meet selection criteria. Parcels identified in Samples 1 and 2 were tested independently in Sample 3.

Using this approach in each sample, we modified our procedure upon discovering that parcels in all networks of interest in Sample 3 (HCP) did not exhibit edges that were significantly positively associated with conscientiousness in 8 or more iterations. We instead combined parcels that met the aforementioned criteria from Samples 1 and 2 to create a larger pool of candidate parcels for each network. Associations between conscientiousness and efficiency of parcels in this pool were then tested independently in Sample 3. This procedure ensured that we could replicate findings from Samples 1 and 2 in a much larger sample, with complete statistical independence from any prior analyses in that sample.

To further refine the search space of influential parcels, we computed the set of combinations of parcels containing a minimum of 3 parcels each to run through the OMST procedure, since a network of 3 nodes is the smallest network for which more than one spanning tree exists (i.e. the smallest fully-connected network with no loops and with potential variability in the configuration of edges). Although these combinations of parcels are treated as “networks” in the graph-theory sense, we use the term “functional ensembles” to describe them from here on, to avoid confusion with our use of “network” to refer to established large-scale brain networks such as the SVAN and its subnetworks. The largest functional ensemble for a given network of interest was defined as containing all parcels meeting the criteria from the permutation-selection procedure in Samples 1 and 2. The largest functional ensembles contained 14 parcels in the SVAN, 5 parcels in subnetwork A, and 9 parcels in subnetwork B. To assess the network-wide functional connectivity of these parcels, we computed measures of efficiency for all functional ensembles.

Partial correlations were computed between conscientiousness and the efficiency of each network or functional ensemble of interest, controlling for age, sex, intelligence, head motion, and efficiency of two other major neural networks (computed using OMST-filtered matrices in the same manner as for the SVAN), the frontoparietal control network (FPCN) and default network (DN) (Yeo et al., 2011). Efficiency values for FPCN and DN were included as covariates for two reasons: First, these networks, like the SVAN, have extensive nodes in prefrontal cortex, and they have additional nodes adjacent to the SVAN throughout the brain. Controlling for their efficiency allows a test of discriminant validity, helping to ensure that any detected associations with conscientiousness are specific to the SVAN, per our hypothesis. Second, a general tendency exists in resting state fMRI for all brain regions and networks to exhibit positive correlations in their activity over time, which may be substantive or artifactual or some combination of both (Rueter et al., 2018). Controlling for the variance in two other large networks removes this general, possibly artifactual variance that the SVAN shares with other parts of the brain.

Intelligence was incorporated into analyses to account for both its modest negative association with conscientiousness and its potentially confounding associations with patterns of functional connectivity in the networks relevant to our hypotheses (Cole et al., 2012; Finn et al., 2015). Removing intelligence as a covariate did not produce appreciable differences in associations between conscientiousness and efficiency values of the networks of interest.

All tests of association between conscientiousness and efficiency of sets of parcels within a particular network were corrected for Type I error using positive false discovery rate (pFDR; Storey, 2002; 2003). To evaluate the significance of each functional ensemble while accounting for the unique distributions of effect sizes in each sample, we utilized the algorithm described by Storey and Tibshirani (2003) to estimate a *q*-value for each functional ensemble, the pFDR analogue of the *p*-value.

### Data/Code Availability

Data from Sample 1 are not able to be shared through open access because participants agreed during the informed consent procedure that their data would not be shared beyond the research team. Data from Sample 3 are available from the Human Connectome Project’s website: https://www.humanconnectome.org/study/hcp-young-adult. Functional connectivity matrices from Sample 2 and scripts used in group-level analyses are available in an Open Science Framework repository: https://osf.io/zju2s/?view_only=8521207b2af540b9bab3b04033744f39

## Results

Descriptive statistics for personality questionnaire measures for all samples are reported in Table 1. Partial correlations between conscientiousness and efficiency of the SVAN, its subnetworks, and their largest functional ensembles are presented in Table 2. All parcels identified in the permutation selection procedure that were used to create combinations of functional ensembles for each network of interest are reported in the online supplement.

**Table 1.** Descriptive statistics for personality measures.

|  | *M* | *SD* | Skew |
| --- | --- | --- | --- |
| Sample 1 (*N* = 244) |  |  |  |
| Self-report BFAS Conscientiousness | 3.39 | .61 | -.20 |
| Self-report BFI Conscientiousness | 3.75 | .63 | -.50 |
| Peer-report BFAS Conscientiousness | 3.42 | .55 | -.17 |
| Peer-report BFI Conscientiousness | 3.66 | .45 | -.57 |
| Composite Conscientiousness | 3.55 | .50 | -.36 |
| Sample 2 (*N* = 239) |  |  |  |
| BFAS Conscientiousness | 3.49 | .51 | -.23 |
| NEO-FFI Conscientiousness | 2.76 | .57 | -.36 |
| Combined Conscientiousness | 3.31 | .61 | -.45 |
| Sample 3 (*N* = 1000) |  |  |  |
| NEO-FFI Conscientiousness | 2.87 | .49 | -.36 |

*Note.* BFAS = Big Five Aspect Scales, BFI = Big Five Inventory, NEO-FFI = NEO Five Factor
Inventory

**Table 2.** Partial correlations between conscientiousness and efficiency of whole networks and largest functional ensembles

| Network | Sample 1<br>(*N* = 244) | Sample 2<br>(*N* = 239) | Sample 3<br>(*N* = 1000) |
| --- | --- | --- | --- |
| SVAN |  |  |  |
| All Parcels | -.01 | .00 | .05* |
| Largest Functional Ensemble | .14** | .17** | .07* |
| Subnetwork A |  |  |  |
| All Parcels | .01 | -.07* | .04 |
| Largest Functional Ensemble | .17** | .06* | .02 |
| Subnetwork B |  |  |  |
| All Parcels | .12* | .17* | .04 |
| Largest Functional Ensemble | .13* | .12* | .02 |

*Note.* Associations controlling for the age, sex, intelligence, head motion, and efficiency of the frontoparietal control and default networks. \* $q < .05$ , \*\* $q < .01$ . SVAN = salience/ventral attention network. Largest functional ensemble refers to the set of all parcels identified during the permutation selection procedure.

### SVAN

The efficiency of the full SVAN, containing all parcels in both subnetworks, was positively associated with conscientiousness in Sample 3, but not Samples 1 and 2. The largest functional ensemble of the SVAN was characterized by a total of 14 parcels located in regions of the left parietal operculum and dorsolateral PFC, right superior frontal gyrus and ventrolateral PFC, and bilateral dACC and insula (Figure 3). The names, indices from Schaefer et al.’s (2018) parcellation scheme (available online at https://github.com/ThomasYeoLab/CBIG/blob/master/stable_projects/brain_parcellation/Schaefe r2018_LocalGlobal/Parcellations/MNI/Centroid_coordinates/Schaefer2018_400Parcels_17Netw orks_order_FSLMNI152_1mm.Centroid_RAS.csv), and centroid coordinates for all parcels in the largest functional ensemble of the SVAN are reported in Table 3. The efficiency of this functional ensemble was significantly associated with conscientiousness across all samples.

**Figure 3.**
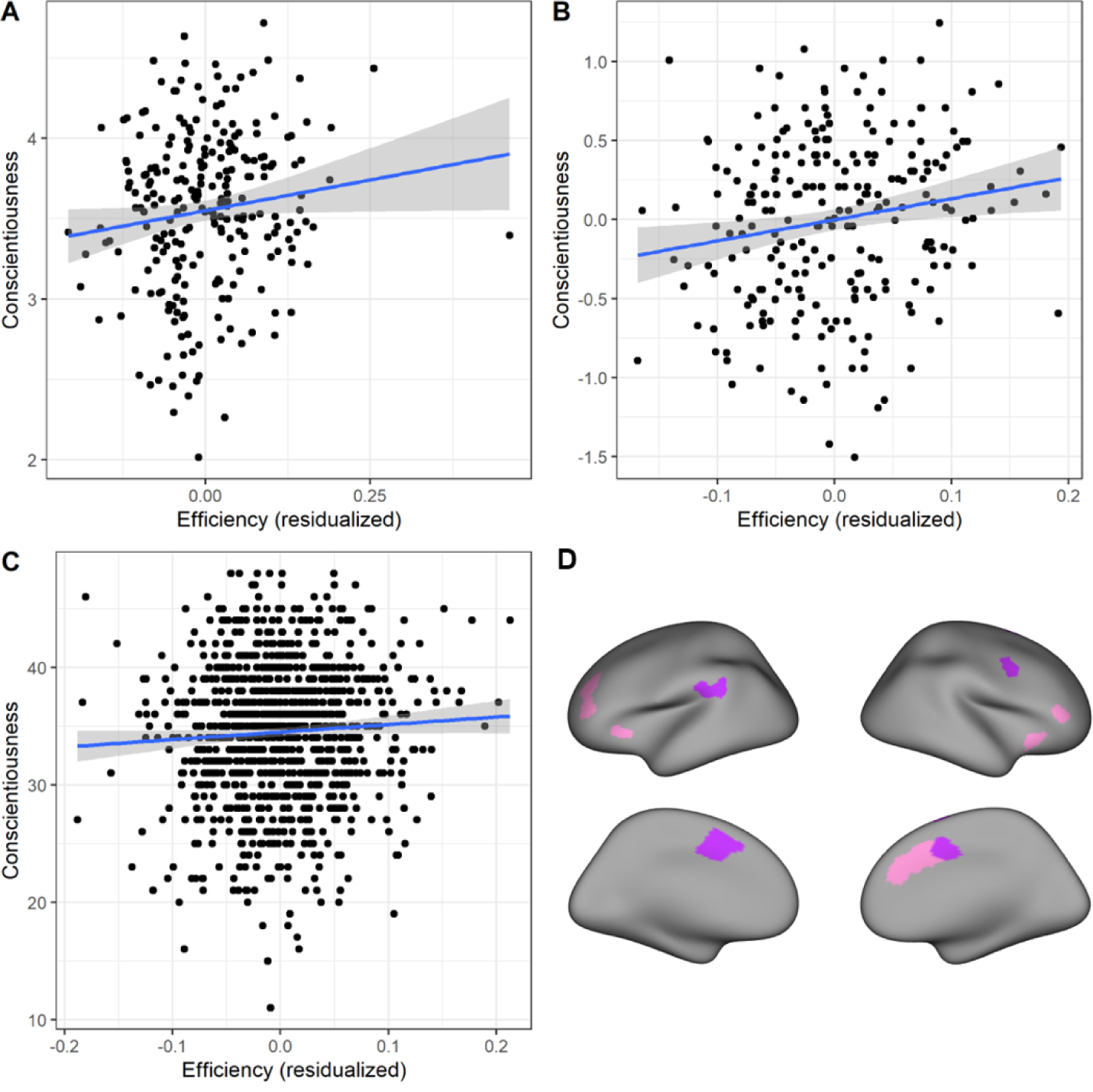
Scatterplots of significant associations between efficiency of the largest functional ensemble of the SVAN with conscientiousness. **A** = Sample 1, **B** = Sample 2, **C** = Sample 3. Associations control for the effects of age, sex, intelligence, and efficiency of the frontoparietal control and default networks. The removal of the outlier in Sample 1 did not appreciably alter the effect, which remained significant (*r* = .11, *q* < .05). **D** = parcels in the largest functional ensemble of the SVAN. SVAN = salience/ventral attention network. Purple = subnetwork A. Pink = subnetwork B.

**Table 3.** MNI152 1mm RAS coordinates for centroids of parcels mapped to the largest functional ensemble of the SVAN

| Parcel Index | Parcel Name | R | A | S |
| --- | --- | --- | --- | --- |
| 86 | LH_SalVentAttnA_ParOper_1 | -55 | -32 | 22 |
| 87 | LH_SalVentAttnA_ParOper_2 | -58 | -44 | 27 |
| 98 | LH_SalVentAttnA_FrMed_1 | -7 | 0 | 57 |
| 99 | LH_SalVentAttnA_FrMed_2 | -5 | 9 | 48 |
| 101 | LH_SalVentAttnB_PFCI_1 | -38 | 49 | 11 |
| 102 | LH_SalVentAttnB_PFCI_2 | -29 | 43 | 30 |
| 105 | LH_SalVentAttnB_Ins_2 | -33 | 25 | -1 |
| 288 | RH_SalVentAttnA_PrC_1 | 51 | 3 | 41 |
| 296 | RH_SalVentAttnA_FrMed_1 | 7 | 2 | 43 |
| 303 | RH_SalVentAttnA_FrMed_4 | 16 | 7 | 69 |
| 305 | RH_SalVentAttnB_PFCIv_1 | 49 | 40 | 5 |
| 309 | RH_SalVentAttnB_Ins_1 | 34 | 21 | -8 |
| 311 | RH_SalVentAttnB_PFCmp_1 | 8 | 35 | 25 |
| 312 | RH_SalVentAttnB_PFCmp_2 | 7 | 19 | 35 |

*Note.* LH = left hemisphere, RH = right hemisphere

Additionally, conscientiousness was positively associated with all combinations of smaller functional ensembles in Sample 2. Similar associations were found in Samples 1 and 3, where conscientiousness was positively associated with 15,616 smaller functional ensembles (96% of all combinations) in Sample 1 and 11,552 smaller functional ensembles (71% of all combinations) in Sample 3, after controlling pFDR (*q* <.05).

### Subnetwork A

In tests of association with subnetwork A of the SVAN, conscientiousness was negatively associated with the efficiency among all parcels in the full subnetwork in Sample 2. However, in Samples 1 and 2, conscientiousness was positively associated with the largest functional ensemble, containing a set of 5 parcels within the left parietal operculum, and the bilateral medial frontal and parietal cortex. Among the combinations of smaller functional ensembles derived from these 5 parcels, conscientiousness was associated with all of these functional ensembles in Sample 1 after pFDR correction for multiple comparisons (*q* < .05), but these associations were not replicated in Samples 2 and 3.

### Subnetwork B

In tests of association with subnetwork B of the SVAN, conscientiousness was significantly associated with the efficiency of all parcels in Samples 1 and 2, but not in Sample 3. Additionally, conscientiousness was significantly associated with the largest functional ensemble in Samples 1 and 2. This functional ensemble contained 9 parcels dispersed throughout the left and right dorsolateral PFC, insula, and dorsomedial PFC. Among the combinations of smaller functional ensembles created using these parcels, conscientiousness was positively associated with 229 functional ensembles (49% of all combinations) in Sample 1 and 164 functional ensembles (35% of all combinations) in Sample 2, after controlling pFDR (*q* < .05).

### Discriminant Validity

To assess the degree to which the association of conscientiousness with network efficiency is specific to the SVAN, we repeated the permutation-selection approach using the parcels of the FPCN and the DN. The largest functional ensemble of the FPCN contained 7 parcels in the bilateral dorsolateral PFC and right anterior cingulate cortex that met the criteria from the permutation-selection procedure in Samples 1 and 2. For this network, efficiency values were computed from OMST-filtered graphs in the same manner as for the SVAN, using all parcels first and then all combinations of functional ensembles using the 7 parcels. We computed partial correlations between conscientiousness and efficiency of the full network, as well as efficiency of all combinations of functional ensembles from these 7 parcels, controlling for the effects of age, sex, intelligence, head motion, and efficiency of all parcels in the SVAN and DN.

This process was then repeated for the DN. The largest functional ensemble of the DN contained 43 parcels which met the criteria from the permutation-selection procedure in Samples 1 and 2. Considering the enormous number of possible combinations of parcels that could be calculated as functional ensembles from this large set of parcels, we limited our analyses to the efficiency values for the OMST-filtered graphs of the full network, the largest functional ensemble, and all combinations of 42, 41, and 40 parcels, in order to limit combinatorial explosion. Although this subset of combinations does not sample all combinations of functional ensembles of the DN, our approach yielded a number of tests roughly comparable to the number conducted for the SVAN, making it a reasonable comparison. Partial correlations controlled for the effects of age, sex, intelligence, head motion and the efficiency of all parcels in the SVAN and FPCN. Parcels in the FPCN and DN used in the combinations of functional ensembles are reported in the online supplement.

Associations between conscientiousness and the full networks and largest functional ensembles of the FPCN and DN are reported in Table 4. Across all samples, conscientiousness was not significantly associated with the efficiency among all parcels of the FPCN. In Samples 1 and 2, conscientiousness was associated with efficiency among the parcels of the largest functional ensemble after controlling pFDR (*q* < .05). Additionally, conscientiousness was significantly associated with the efficiency of 64 functional ensembles (64% of combinations) in Sample 1 after controlling pFDR (*q* < .05), and the efficiency of all functional ensembles in Sample 2 after controlling pFDR (*q* < .05). However, none of these associations were replicated in Sample 3, which is the crucial test because Sample 3 was not involved in selecting the parcels in the first place. In tests of DN efficiency, conscientiousness was not significantly associated with the efficiency among all parcels of the DN in all samples after controlling pFDR (*q* < .05). Conscientiousness was significantly associated with the efficiency of all functional ensembles in Sample 2 after controlling pFDR (*q* < .05), but these associations were not replicated in Samples 1 and 3. The lack of replicable associations between conscientiousness and efficiency in the DN and FPCN provides evidence of discriminant validity, demonstrating that associations with the SVAN are probably not attributable to factors like network size or general characteristics of brain network integration.

**Table 4.** Partial correlations between conscientiousness and efficiency of whole networks and largest functional ensembles in FPCN and DN

| Network | Sample 1<br>(*N* = 244) | Sample 2<br>(*N* = 239) | Sample 3<br>(*N* = 1000) |
| --- | --- | --- | --- |
| FPCN |  |  |  |
| All Parcels | .03 | -.04 | -.01 |
| Largest Functional Ensemble | .14* | .17** | .03 |
| DN |  |  |  |
| All Parcels | .00 | .08 | .03 |
| Largest Functional Ensemble | .07 | .20** | .02 |

*Note.* Associations controlling for the age, sex, intelligence, head motion, and efficiency of the SVAN. Associations with FPCN efficiency control for whole DN efficiency, and vice versa. \* $q < .05$ , \*\* $q < .01$ . FPCN = frontoparietal control network. DN = default network. Largest functional ensemble refers to the set of all parcels identified during the permutation selection procedure.

To further assess discriminant validity, we extended our analyses to determine whether these associations with SVAN were exclusive to conscientiousness relative to the other Big Five dimensions. For each sample, we repeated our analyses including the remaining Big Five traits as simultaneous predictors in tests of association between conscientiousness and efficiency of the full SVAN and combinations of functional ensembles. Partial correlations between each of the Big Five and efficiency of all parcels in the SVAN and the largest functional ensemble are reported in Table 5. Among associations between traits other than conscientiousness, efficiency values of the full SVAN and its largest functional ensemble in Sample 1 were significantly negatively associated with neuroticism after controlling pFDR (*q* < .05), but this association was not replicated in Samples 2 and 3. Interestingly, conscientiousness was not significantly associated with efficiency of the full SVAN or its largest functional ensemble in Samples 1 and 2 after controlling for the remaining Big Five and controlling pFDR (*q* < .05), but conscientiousness in Sample 3 remained significantly positively associated with efficiency in the full SVAN and 14,330 functional ensembles (88% of all combinations), including the largest functional ensemble, after controlling for the remaining Big Five and controlling pFDR (*q* < .05). Although the partial association between conscientiousness and efficiency of the SVAN was statistically significant only in Sample 3, the magnitude of the effect in Sample 1 mirrors that of Sample 3, suggesting the possibility that these results in Sample 1 may be indicative of Type II error as a result of lower statistical power in a smaller sample. Regardless, conscientiousness was the only trait among the Big Five to consistently exhibit positive associations with efficiency of the largest functional ensemble of the SVAN across samples, providing further evidence of discriminant validity.

**Table 5.** Partial correlations between the Big Five and efficiency of the SVAN and its largest functional ensemble, controlling for other Big Five dimensions.

| SVAN | C | E | A | N | O/I |
| --- | --- | --- | --- | --- | --- |
| Sample 1 ($N = 244$) | | | | | |
| All Parcels | -.04 | .05 | -.06 | -.08* | .04 |
| Largest Functional Ensemble | .07 | .01 | -.05 | -.11* | .00 |
| Sample 2 ($N = 239$) | | | | | |
| All Parcels | .00 | .00 | -.02 | .02 | .00 |
| Largest Functional Ensemble | .01 | -.01 | -.01 | .04 | .00 |
| Sample 3 ($N = 1000$) | | | | | |
| All Parcels | .07* | .04 | -.05 | .05 | .02 |
| Largest Functional Ensemble | .07* | .05 | -.04 | .04 | .04 |

*Note.* Associations controlling for the age, sex, intelligence, head motion, and efficiency of the frontoparietal control and default networks. \* $q < .05$ , \*\* $q < .01$ . SVAN = salience/ventral attention network. C = Conscientiousness, E = Extraversion, A = Agreeableness, N = Neuroticism, O/I = Openness/Intellect. Largest functional ensemble refers to the set of all parcels identified during the permutation selection procedure.

## Discussion

This research presents evidence of an association of conscientiousness with the functional connectivity of a set of parcels within the SVAN in three samples. These parcels, located in regions of medial and lateral PFC, insula, and parietal operculum, are widely dispersed throughout many nodes in both subnetworks of the SVAN identified by the 17-network atlas of Yeo et al. (2011). Considering the nature of our parcel selection procedure in constructing subsets of potentially relevant parcels, the magnitude of these associations may be overestimated in the first two samples. However, because the much larger third sample (HCP) did not contribute any parcels to this procedure, the significant effect in HCP is noteworthy as a fully independent and unbiased replication.

The effect is small, but this should not be surprising given that participants were merely resting rather than being required to engage actively in some task involving goal prioritization. In personality neuroscience generally, we should expect the functional properties of networks to be more strongly associated with relevant traits when those networks are actively carrying out the psychological functions corresponding to the trait in question. The magnitude of the correlation between conscientiousness and the largest functional ensemble of the SVAN in Sample 3 is consistent with research on the range of effect sizes for associations between behavioral individual differences and resting brain function, making the large size of Sample 3 important for our ability to test this effect (Marek et al., 2022).

Associations between conscientiousness and the SVAN are consistent with previous research describing the network’s functional roles. In its initial conception, the ventral attention network was largely right-lateralized and included the TPJ and regions of ventral PFC, and it has been described as responsible for monitoring and redirecting attentional processes as a response to new and potentially salient cues from the environment (Corbetta & Shulman, 2002; Fox et al., 2006; Vossel et al., 2014). Researchers identifying a functionally similar network called the salience network, consisting of the dorsomedial PFC, dACC, insula, and select regions of the dorsolateral PFC, have described this network as sharing a similar function to the ventral attention network, but with greater emphasis on integration of information from a variety of sources to discern the relevance of incoming stimuli to one’s motivation and goals (Menon & Uddin, 2010; Seeley et al., 2007; Uddin, 2015). Although previous research has sometimes characterized the ventral attention and salience networks as independent functional systems (Power et al., 2011; Cole et al., 2013; Baker et al., 2014), their anatomical and functional overlap has led to a growing consensus that they refer to a single large network with identifiable subnetworks (Uddin et al., 2019).

The parcels in which efficiency was consistently associated with conscientiousness were spread across both SVAN subnetworks, suggesting that conscientious individuals may be characterized by effective integration of nodes relevant to all functions of the SVAN. This efficiency may permit a greater capability to identify and manipulate motivationally salient information in the context of multiple goals of varying duration, importance, and abstraction.

These findings also have important implications in the context of evidence supporting the involvement of the SVAN in modulating the activity of the DN and FPCN (Kucyi et al., 2017; Zhou et al., 2018). Through this interpretation, our results suggest that individual differences in conscientiousness may be contingent on the differences in the adeptness of the SVAN in managing the dynamic tension between these two networks. This particular role of the SVAN coheres well with the notion of goal prioritization as the balance between manipulating both the internal representations of various goal states and information in working memory used to inform the progress of these goals and direct immediate action related to them.

Further, this interpretation is consistent with much of the literature describing the functional and structural properties of regions where the parcels in question are located, particularly the dorsolateral PFC, insula, and dorsomedial PFC. Previous literature has described associations between functional connectivity among these regions and a variety of higher-order self-regulatory processes conceptually related to conscientiousness (Cieslik et al., 2013; Lynn et al., 2014; Taren et al., 2011). Several structural MRI studies have found conscientiousness to be associated with structural variables such as cortical thickness in regions of dorsolateral PFC corresponding reasonably well to particular parcels identified in the present research (Bjørnebekk et al., 2013; Gao et al., 2021; Owens et al., 2019). These regions of the dorsolateral PFC may be particularly responsible for maintaining the representations that regulate the pursuit of active goals and suppress distractions in relation to salient information identified through other SVAN regions such as the insula (Dosenbach et al., 2008; Rueter et al., 2018). Thus, our results describe replicable associations of conscientiousness with functional integration in the SVAN in a manner that seems consonant with previous literature describing the neural correlates of well-regulated goal pursuit.

### Comparison with Previous Research

Our findings serve as an important clarification and extension of findings by Rueter et al. (2018), supporting the theory that the SVAN is a key neurological substrate of individual differences in conscientiousness. However, our methods allowed us to make a more precise and generalizable test of the hypothesis that functional connectivity in the SVAN is positively associated with conscientiousness. Because Rueter et al. (2018) relied on ICA to identify intrinsic connectivity networks (ICNs) that overlapped with the SVAN more than with other canonical networks, the boundaries of the ICNs did not correspond particularly well to the SVAN’s boundaries. In our supplemental materials, we report analyses in which we tried to replicate, as closely as possible using our parcellation, the spatial extent of the ICN that Rueter et al. (2018) found to be associated with conscientiousness. This included parcels from several other networks in addition to the SVAN. In the same sample that they used (our Sample 1), we found that efficiency among the parcels corresponding to their ICN was indeed significantly associated with conscientiousness, thus showing the reproducibility of their results using our methods. However, this finding did not replicate across samples. This pattern of results suggests the value of working with standard atlases and individualized parcellation, rather than ICA, which produces unique and nonstandard components in every sample. Results based on standard atlases should be more generalizable.

Although the association of conscientiousness with Rueter et al.’s (2018) ICN was not replicated in all samples, the set of parcels in the SVAN that did exhibit a replicable association shared several parcels with that ICN, including parcels in the left dorsolateral PFC, right insula, and right dorsomedial PFC. This corroborates the observation by Rueter et al. (2018) that their ICN shared considerable overlap with subnetwork B of the SVAN. Thus, the basic conclusions of Rueter et al. regarding the importance of the SVAN for conscientiousness appears to have been correct, even though their use of ICA rendered the exact brain regions they identified less generalizable than the results of the current study.

Much of the previous literature investigating individual differences in functional connectivity has used a similar procedure to Rueter et al. (2018): ICA followed by dual regression (Poppe et al., 2013). ICA is an exploratory procedure that identifies patterns of covariance among voxels in the sample in question, leading to a set of components at the group level that are interpreted as intrinsic connectivity networks (ICNs). These ICNs are then individualized by dual regression, in which the group level map is applied spatially to each subject, then the aggregate timeseries of the voxels in a given ICN is regressed on the subject-specific map, thereby adjusting the location of the ICN to be unique to each subject.

There are at least three serious limitations to this procedure. First, dual regression rests on the obviously problematic assumption that the aggregate timeseries of what is known to be the wrong collection of voxels can be used to identify the right collection of voxels. The more a given subject’s ICN is in a different location from the group level ICN, the more correction is needed, yet the less accurate the correction will be given that the starting values will be farther from the correct values. Second, because ICA is an exploratory procedure, its results are different in every sample, and it can be difficult or impossible to match ICNs across samples. (We found this to be the case when we compared results from Rueter et al.’s ICA with the ICA results publicly available for HCP.) Third, group level ICNs often do not align well with network boundaries in standard atlases, making tests of hypotheses regarding canonical networks ambiguous.

Use of individualized parcellation with a standard atlas was one way we improved on previous research, and use of OMSTs to assess functional connectivity is another. OMSTs capture meaningful individual variability in functional topology among parcels by modeling associations among them in a manner that maximizes global-cost efficiency, consequently bypassing both the need to impose an arbitrary threshold and the retention of excess noise in graph theoretical measures. By simultaneously modeling efficient information transfer and sparsity using OMSTs, our approach effectively characterizes brain connectivity as high-performing and adaptable small-world networks with modular and hierarchical properties, which corresponds well to what is known about actual brain function (Bassett and Bullmore, 2006; Mengistu et al., 2016). The present research incorporates many practices that have been shown in recent work to improve test-retest reliability among resting-state fMRI findings, including the use of the Schaefer atlas (Schaefer et al., 2018), weighted graphs determined by product-moment correlations, and graph filtering through OMSTs (Luppi et al., in review; Luppi & Stamatakis, 2021).

### Limitations

Despite the advantages of our methods, there are limitations to consider regarding the current research. Although we identified a set of parcels for which associations with conscientiousness were replicable in a large, independent sample, we also saw some variability across samples in associations between conscientiousness and the efficiency of various networks and functional ensembles. This variability could be related to variability in participant characteristics, scanning parameters, and/or personality measures. For example, quality of measurement might explain why Sample 1, with multiple self- and peer-report measures of personality, demonstrated stronger associations between neural variables and conscientiousness than Sample 3, which had only a single, relatively brief self-report measure of personality.

Additionally, the permutation-based parcel selection procedure used to identify subnetworks for each of the canonical networks of interest used a cutoff of parcels appearing in 80% of iterations. This cutoff was chosen to permit a suitably sized pool of candidate parcels while also affording manageable computational intensity during the calculation of functional ensemble combinations. This somewhat arbitrary threshold defined the size of the largest functional ensemble for each network, and a different threshold might have yielded different results.

## Conclusion

Using resting state fMRI, in conjunction with canonical networks of individualized parcels and graph theoretical measures, we identified a set of regions in the SVAN, including nodes in the anterior insula, dorsolateral PFC, parietal operculum, and dACC, in which network efficiency was a replicable neural correlate of conscientiousness across three samples. These results are consistent with a theory that the ability to prioritize goals effectively underlies conscientiousness and relies primarily on the SVAN (Allen & DeYoung, 2017; Rueter et al., 2018). Our findings emphasize the importance of several existing and emerging practices in personality neuroscience (DeYoung et al., in press), including the use of theory-driven analyses and large sample sizes, as well as the use of individualized parcellation to capture variability in the cortical location of canonical neural networks. Although this is basic research, it may eventually contribute to the development of novel interventions for mental health problems involving impulsivity and disinhibition, the dimension of psychopathology corresponding to maladaptive low conscientiousness.

## Acknowledgements

We would like to thank Amanda Rueter, Katherine Scotting, and Ranee Flores for their feedback and input in this work. Data collection was supported by grants from the National Institute on Drug Abuse (R03DA029177-01A1) and the National Science Foundation (SES- 1061817) to Colin DeYoung and Aldo Rustichini, and from the John Templeton Foundation (22156) to Rex Jung. We also thank the collectors and curators of the data from the Human Connectome Project.

## Online Supplement

### Parcels of Functional Ensembles

**Table A1.**
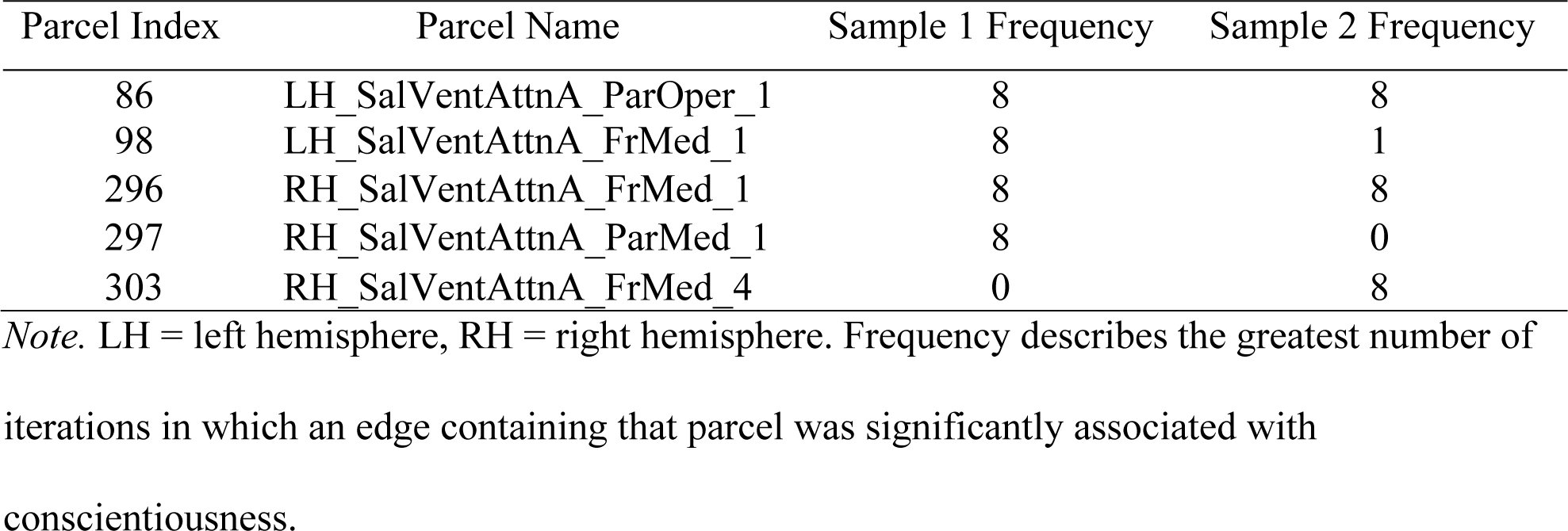
Parcels used to define functional ensembles in subnetwork A

**Table A2.** Parcels used to define functional ensembles of subnetwork B

| Parcel Index | Parcel Name | Sample 1 Frequency | Sample 2 Frequency |
| --- | --- | --- | --- |
| 102 | LH_SalVentAttnB_PFCI_2 | 10 | 6 |
| 104 | LH_SalVentAttnB_Ins_1 | 2 | 8 |
| 106 | LH_SalVentAttnB_Ins_3 | 0 | 8 |
| 108 | LH_SalVentAttnB_PFCmp_1 | 9 | 1 |
| 305 | RH_SalVentAttnB_PFCIv_1 | 1 | 10 |
| 306 | RH_SalVentAttnB_PFCIv_1 | 9 | 3 |
| 309 | RH_SalVentAttnB_Ins_1 | 0 | 10 |
| 311 | RH_SalVentAttnB_PFCmp_1 | 10 | 7 |
| 312 | RH_SalVentAttnB_PFCmp_2 | 9 | 6 |

*Note.* LH = left hemisphere, RH = right hemisphere. Frequency describes the greatest number of iterations in which an edge containing that parcel was significantly associated with conscientiousness.

**Table A3.** Parcels used to define subnetworks of the SVAN

| Parcel Index | Parcel Name | Sample 1 Frequency | Sample 2 Frequency |
| --- | --- | --- | --- |
| 86 | LH_SalVentAttnA_ParOper_1 | 8 | 8 |
| 87 | LH_SalVentAttnA_ParOper_2 | 0 | 10 |
| 98 | LH_SalVentAttnA_FrMed_1 | 10 | 4 |
| 99 | LH_SalVentAttnA_FrMed_2 | 9 | 3 |
| 101 | LH_SalVentAttnB_PFCI_1 | 3 | 10 |
| 102 | LH_SalVentAttnB_PFCI_2 | 10 | 7 |
| 105 | LH_SalVentAttnB_Ins_2 | 1 | 10 |
| 288 | RH_SalVentAttnA_PrC_1 | 1 | 10 |
| 296 | RH_SalVentAttnA_FrMed_1 | 4 | 10 |
| 303 | RH_SalVentAttnA_FrMed_4 | 2 | 8 |
| 305 | RH_SalVentAttnB_PFCIv_1 | 0 | 9 |
| 309 | RH_SalVentAttnB_Ins_1 | 0 | 9 |
| 311 | RH_SalVentAttnB_PFCmp_1 | 10 | 10 |
| 312 | RH_SalVentAttnB_PFCmp_2 | 6 | 9 |

*Note.* LH = left hemisphere, RH = right hemisphere. Frequency describes the greatest number of
iterations in which an edge containing that parcel was significantly associated with conscientiousness.

**Table A4.** Parcels used to define subnetworks of the FPCN

| Parcel Index | Parcel Name | Sample 1 Frequency | Sample 2 Frequency |
| --- | --- | --- | --- |
| 128 | LH_ContA_PFCIv_1 | 9 | 8 |
| 140 | LH_ContB_PFCIv_1 | 4 | 8 |
| 330 | RH_ContA_PFCI_1 | 0 | 10 |
| 334 | RH_ContA_PFCI_5 | 0 | 10 |
| 335 | RH_ContA_Cingm_1 | 9 | 0 |
| 346 | RH_ContB_PFCIv_1 | 0 | 10 |
| 347 | RH_ContB_PFCIv_2 | 0 | 10 |

*Note.* LH = left hemisphere, RH = right hemisphere. Frequency describes the greatest number of
iterations in which an edge containing that parcel was significantly associated with
conscientiousness.

**Table A5.** Parcels used to define subnetworks of the DN

| Parcel Index | Parcel Name | Sample 1 Frequency | Sample 2 Frequency |
| --- | --- | --- | --- |
| 150 | LH_DefaultA_IPL_2 | 1 | 10 |
| 151 | LH_DefaultA_PFCd_1 | 1 | 10 |
| 152 | LH_DefaultA_PFCd_2 | 2 | 10 |
| 155 | LH_DefaultA_pCunPCC_2 | 3 | 10 |
| 161 | LH_DefaultA_PFCm_1 | 1 | 8 |
| 163 | LH_DefaultA_PFCm_3 | 8 | 0 |
| 164 | LH_DefaultA_PFCm_4 | 8 | 0 |
| 165 | LH_DefaultA_PFCm_5 | 0 | 10 |
| 166 | LH_DefaultA_PFCm_6 | 6 | 8 |
| 167 | LH_DefaultB_Temp_1 | 4 | 10 |
| 170 | LH_DefaultB_Temp_4 | 8 | 10 |
| 171 | LH_DefaultB_Temp_5 | 1 | 10 |
| 174 | LH_DefaultB_IPL_2 | 3 | 10 |
| 176 | LH_DefaultB_PFCd_2 | 2 | 10 |
| 177 | LH_DefaultB_PFCd_3 | 8 | 9 |
| 178 | LH_DefaultB_PFCd_4 | 10 | 10 |
| 179 | LH_DefaultB_PFCd_5 | 7 | 8 |
| 182 | LH_DefaultB_PFCl_2 | 3 | 10 |
| 183 | LH_DefaultB_PFCv_1 | 0 | 10 |
| 185 | LH_DefaultB_PFCv_3 | 6 | 10 |
| 186 | LH_DefaultB_PFCv_4 | 10 | 8 |
| 187 | LH_DefaultB_PFCv_5 | 10 | 10 |
| 189 | LH_DefaultC_Rsp_1 | 0 | 8 |
| 191 | LH_DefaultC_Rsp_3 | 1 | 10 |
| 358 | RH_DefaultA_Temp_1 | 0 | 8 |
| 360 | RH_DefaultA_IPL_2 | 0 | 10 |
| 364 | RH_DefaultA_pCunPCC_2 | 1 | 10 |
| 370 | RH_DefaultA_PFCm_3 | 10 | 7 |
| 371 | RH_DefaultA_PFCm_4 | 8 | 8 |
| 372 | RH_DefaultA_PFCm_5 | 1 | 10 |
| 373 | RH_DefaultA_PFCm_6 | 10 | 7 |
| 374 | RH_DefaultB_Temp_1 | 0 | 10 |
| 375 | RH_DefaultB_Temp_2 | 3 | 8 |
| 376 | RH_DefaultB_AntTemp_1 | 3 | 10 |
| 377 | RH_DefaultB_PFCd_1 | 7 | 10 |
| 378 | RH_DefaultB_PFCd_2 | 1 | 9 |
| 379 | RH_DefaultB_PFCd_3 | 8 | 3 |
| 380 | RH_DefaultB_PFCd_4 | 0 | 10 |
| 381 | RH_DefaultB_PFCd_5 | 4 | 8 |
| 382 | RH_DefaultB_PFCv_1 | 6 | 9 |
| 383 | RH_DefaultB_PFCv_2 | 10 | 10 |
| 384 | RH_DefaultB_PFCv_3 | 4 | 9 |
| 385 | RH_DefaultC_IPL_1 | 1 | 8 |

*Note.* LH = left hemisphere, RH = right hemisphere. Frequency describes the greatest number of iterations in which an edge containing that parcel was significantly associated with conscientiousness.

**Table A6.** MNI152 1mm RAS coordinates for centroids of parcels mapped to ICN1

| Parcel Index | Parcel Name | R | A | S |
| --- | --- | --- | --- | --- |
| 102 | LH_SalVentAttnB_PFCI_2 | -29 | 43 | 30 |
| 104 | LH_SalVentAttnB_Ins_1 | -34 | 16 | -8 |
| 106 | LH_SalVentAttnB_Ins_3 | -33 | 25 | -1 |
| 108 | LH_SalVentAttnB_PFCmp_1 | -6 | 22 | 31 |
| 133 | LH_ContA_Cingm_1 | -3 | 5 | 29 |
| 143 | LH_ContB_PFCmp_1 | -4 | 28 | 47 |
| 164 | LH_DefaultA_PFCm_4 | -6 | 45 | 6 |
| 166 | LH_DefaultA_PFCm_6 | -5 | 34 | 21 |
| 175 | LH_DefaultB_PFCd_1 | -4 | 51 | 28 |
| 183 | LH_DefaultB_PFCv_1 | -36 | 22 | -16 |
| 307 | RH_SalVentAttnB_PFCI_2 | 25 | 54 | 25 |
| 309 | RH_SalVentAttnB_Ins_1 | 34 | 21 | -8 |
| 311 | RH_SalVentAttnB_PFCmp_1 | 8 | 35 | 25 |
| 312 | RH_SalVentAttnB_PFCmp_2 | 7 | 19 | 35 |
| 335 | RH_ContA_Cingm_1 | 5 | 1 | 30 |
| 350 | RH_ContB_PFCmp_1 | 5 | 28 | 48 |
| 370 | RH_DefaultA_PFCm_3 | 7 | 42 | 4 |
| 373 | RH_DefaultA_PFCm_6 | 6 | 25 | 18 |
| 379 | RH_DefaultB_PFCd_3 | 5 | 44 | 40 |
| 382 | RH_DefaultB_PFCv_1 | 35 | 23 | -18 |

*Note.* LH = left hemisphere, RH = right hemisphere

**Table A7.** Partial correlations between conscientiousness and efficiency of ICN1

| Network | Sample 1<br>(*N* = 244) | Sample 2<br>(*N* = 239) | Sample 3<br>(*N* = 1000) |
| --- | --- | --- | --- |
| ICN1 | .19** | -.07 | .06 |

*Note.* Associations controlling for the age, sex, intelligence, head motion, and efficiency of the frontoparietal control and default networks. \*\* $p < .01$ . ICN1 = intrinsic connectivity network 1.

#### Replication of Findings by Rueter et al. (2018)

In addition to the parcels of the SVAN, we also tested whether parcels corresponding to an intrinsic connectivity network (ICN) previously found by Rueter et al. (2018) to be associated with conscientiousness would yield replicable results across samples. Thus, in addition to having a test of our theoretically derived hypothesis in multiple samples, we could also compare these results to a test of an empirically derived hypothesis, namely that the ICN identified using ICA in Rueter et al.’s analysis might represent a robust correlate of conscientiousness.

To evaluate associations between conscientiousness and the ICN designated as “ICN1” by Rueter et al. (2018), which they found to be most strongly associated with conscientiousness, Schaefer et al.’s (2018) volumetric 400-parcel group-level atlas in MNI space was overlaid with a binarized mask of the voxels within ICN1 at the group level. Any parcel exhibiting 50% or greater overlap with voxels within ICN1 was selected as belonging to this network. This particular cutoff was used to identify parcels exhibiting substantial overlap with ICN1, in an attempt to map the volumetric ICA results of Rueter et al. (2018) to a surface-based parcellation. All parcels meeting this criterion are illustrated using the group-level Schaefer atlas (Schaefer et al., 2018) in Figure A1. Centroid coordinates, names, and atlas indices for all parcels overlapping with ICN1 are reported in Table A6.

Partial correlations were computed between conscientiousness and the efficiency of ICN1 controlling for age, sex, intelligence, head motion, and the efficiency of the FPCN and DN. Because ICN1 contained a selection of parcels belonging to the FPCN and DN, the efficiency covariates were computed using the remaining FPCN and DN parcels outside of ICN1.

In agreement with Rueter et al. (2018), conscientiousness was significantly associated with efficiency among all parcels in ICN1 in the original sample (Sample 1). However, these associations did not replicate in Samples 2 and 3. Associations between conscientiousness and efficiency of ICN1 are described in Table A7.

**Figure A1.**
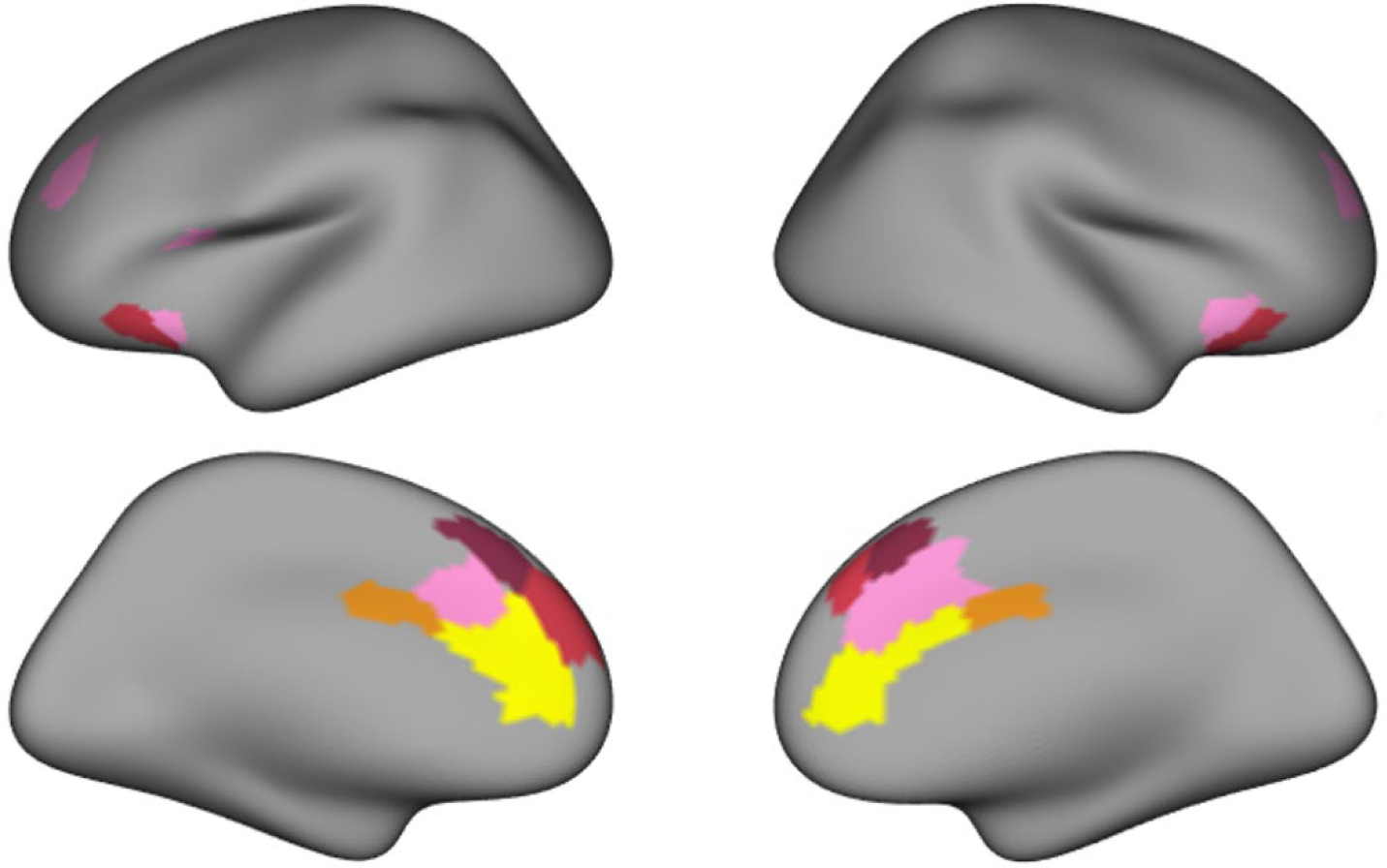
Parcels in ICN1. Colors correspond to 17 functional networks described by Yeo et al. (2011). Yellow = default subnetwork A, Red = default subnetwork B, Orange = frontoparietal control subnetwork A, Maroon = frontoparietal control subnetwork B, Pink = SVAN subnetwork B.

